# Identification of Cyclin L1 as a host factor regulating Hepatitis B Virus replication

**DOI:** 10.1101/2024.10.23.619969

**Authors:** Collins Oduor Owino, Balakrishnan Chakrapani Narmada, Gian Yi Lin, Pauline Poh Kim Aw, Nivrithi Ganesh, Jovi Tan Siying, Marie-Laure Plissonnier, Thangavelu Thangavelu Matan, Niranjan Shirgaonkar, Juan Pablo Bifani, Massimo Levrero, Giridharan Periyasamy, Seng Gee Lim, Ramanuj DasGupta

## Abstract

**Background & Aims:** Understanding the regulatory interactions between Hepatitis B virus (HBV) and human host factors is key to the development of next-generation host-directed antiviral therapies and achieving a functional HBV cure. In this study, we aimed to investigate HBV-induced alterations in host gene expression in primary human hepatocytes (PHH) to identify host-specific factors that are regulated and exploited by the virus for replication and survival.

**Methods:** We performed whole transcriptome sequencing (WTS) of HBV-infected PHH to identify host pathways that could potentially influence the HBV life cycle. RNA-interference-based validation of putative targets to evaluate the function of dysregulated candidate genes resulted in the identification of Cyclin L1 (CCNL1) as a key host factor.

**Results:** RNAi-knockdown of *CCNL1* revealed that it is essential for HBV gene expression, including HBV-surface antigen (HBsAg). Mechanistically, we found that CCNL1 can phosphorylate the C-terminal domain (CTD) of RNA Polymerase II (RNAPII) at serine 2 (S2), likely to regulate HBV transcription. Furthermore, the knockdown of CCNL1 inhibited the binding of total and phospho- (Ser2-Ser5) RNAPII, pan-acetylated H3ac, and H3K27ac to HBV cccDNA, implicating its function in the regulation of cccDNA-dependent viral transcription. Finally, enhanced CCNL1 expression in chronic hepatitis B patients, as compared to those with resolved infection, underscores a functional link between this host factor and CHB.

**Conclusion:** Our data demonstrates that CCNL1 regulates HBV RNA transcription and replication by modulating RNAPII phosphorylation and activity, making it a potential host susceptibility factor for HBV.

**LAY SUMMARY:** Hepatitis B requires human host cell factors and biological processes to establish an efficient infection. Identifying host factors that support and/or restrict HBV infection is essential for understanding the molecular basis of chronic HBV infection and for developing host-targeting anti-HBV drugs. Here, we report that *CCNL1* can serve as a potential host susceptibility factor for HBV, as reduced CCNL1 function results in reduced viral replication and gene expression.

**Graphical Summary:** Graphical summary, highlighting the approach and validation experiments. From whole transcriptomics analysis, we identified known HBV-host factors such as SRPK1, CDK1, NXF1 among others as well as new factors such as CCNL1. Employing various RNAi approaches and different cell models including primary human hepatocytes, we validated the role of CCNL1 during HBV infection cycle. Pull-down experiments followed by ChIP-PCR further showed a reduction in cccDNA-based transcription upon knockdown of CCNL1.

**Graphical abstract:** 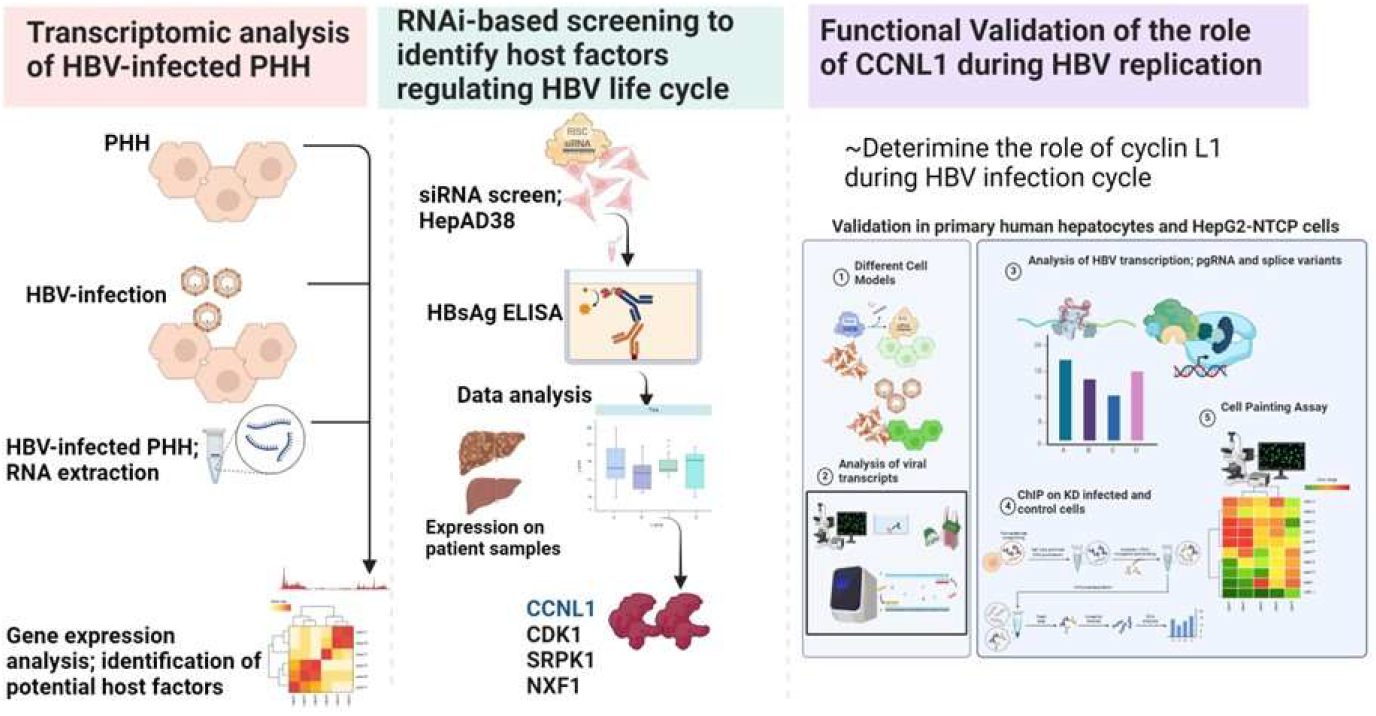

## INTRODUCTION

Chronic Hepatitis B (CHB) infection affects over 250 million people worldwide and can lead to significant liver disease with long-term health consequences including liver damage, failure, and cancer (Lampertico *et al*, 2017; Narmada *et al*, 2024). Clinical management of CHB includes nucleos(t)ide analogs (NAs) and interferon (IFN)-based therapy (Lampertico *et al*, 2017; Narmada *et al*, 2024). While current therapeutic strategies suppress viral replication and reduce the risk of developing adverse liver disease, they cannot completely eliminate the virus or achieve a durable functional cure, as defined by the loss of HBV surface antigen (HBsAg) (Lampertico *et al*, 2017; Zeisel *et al*, 2015; Ghany, 2017; Fanning *et al*, 2019). These limitations call for the implementation of novel strategies to identify the next generation of anti-HBV drug targets and drugs to boost ongoing efforts to eradicate HBV infection.

HBV is a small, partially double-stranded DNA virus belonging to the family Hepadnaviridae that exhibits high tropism for human hepatocytes. HBV encodes seven viral proteins and only one enzyme, DNA polymerase. Therefore, the virus relies heavily on host cell factors to infect and replicate in human hepatocytes. These host factors may either be intrinsic host factors that the virus coopts, or host restriction factors (HRFs), such as innate signalling pathways and defense mechanisms, which the virus must suppress to establish an efficient infection.

Although numerous host factors regulating HBV replication have been described, the importance and mechanism of action of various human cellular factors at different stages of the viral infection cycle remain unclear. For example, ectopic expression of sodium taurocholate (NTCP), the major receptor for HBV, in mouse primary hepatocytes does not render them susceptible to HBV infection (Yan *et al*, 2013), and different HepG2-NTCP clones display different degrees of susceptibility to HBV infection (Iwamoto *et al*, 2014). These observations suggest that other, as yet unknown, host factors are involved in establishing robust and efficient HBV infection. Thus, a deeper probing of host-pathogen interactions is needed to not only expand our knowledge of viral pathogenesis but may also result in the identification of novel anti-HBV therapy that can be used in combination with current therapies to achieve better, and durable treatment outcomes.

In this study, we employed whole-transcriptome sequencing (WTS) to identify HBV-mediated gene perturbations in physiologically relevant primary human hepatocytes (PHH) to identify unique infection-driven host gene expression signatures. These genes were functionally investigated for their potential function as novel host susceptibility factors for HBV replication and transcription. Specifically, we report the identification of Cyclin L1 (CCNL1) as a possible HBV host factor that affects viral transcription. Loss of *CCNL1* function results in a global decrease in HBV gene expression, whereas its ectopic expression leads to increased viral replication. Mechanistically, we demonstrated that *CCNL1* regulates the transcription of nascent HBV RNA by modulating RNAPII activity and the subsequent epigenetic regulation of cccDNA. Finally, we show that CHB patients display a general increase in CCNL1 expression compared to those who have achieved a functional cure. Overall, these results corroborate the physiological role of CCNL1 in the control of HBV replication/transcription and highlight its value as a putative host-directed therapeutic target for the treatment of CHB.

## RESULTS

### Identification of CCNL1 as a host susceptibility factor in HBV infection

Primary human hepatocytes (PHH) offer a physiologically relevant model for identifying and validating human host factors that regulate HBV infection (Schulze-Bergkamen *et al*, 2003; Hu *et al*, 2019; Galle *et al*, 1994). We hypothesized that HBV infection alters the host transcriptome to support its replication and transcription. Therefore, we performed an integrated transcriptomic analysis of HBV-infected PHH combined with RNAi-mediated loss-of-function studies to validate the potential role(s) in the regulation of various aspects of the HBV life cycle (**Fig. 1A**). First, we performed bulk RNA-seq analysis of HBV-mediated transcriptomic changes in PHH to identify host genes that were either up- or down-regulated during HBV infection, compared to mock-infected controls (see volcano plot in **Fig. 1B**). Gene set enrichment analysis (GSEA) of differentially expressed genes 96 h post HBV infection revealed 69 significantly upregulated genes based on a log_2_ fold change>1.5 and p-value<0.01 (**Fig. 1B, C**). Pathway analysis of these genes revealed enrichment of previously described cellular processes associated with HBV infection, including the G2/M checkpoint, solute carriers, and p38 mitogen-activated protein kinase (MAPK) (**Fig. 1C**) (Chang *et al*, 2008; Lamontagne *et al*, 2016). Intriguingly, our study also identified the enrichment of unique pathways that bear the hallmarks of cancer, including EMT, the core matrisome, Aurora B kinase, and VEGFR signalling (**Fig. 1C**). In contrast, pathways that are known to restrict HBV infection, such as hypoxia, were downregulated in our analysis (**Fig. S1A)** (Hallez *et al*, 2019). Additionally, consistent with existing evidence linking HBV infection and reprogramming and/or alterations in host cell metabolism (Masson *et al*, 2017; Li *et al*, 2015), we identified several pathways involved in various metabolic processes that were deregulated upon HBV infection in PHH (**Fig. 1C, Fig. S1B**). Together, these results enabled the characterization of HBV-induced alterations in the host transcriptome of PHH upon infection.

**Figure 1:**
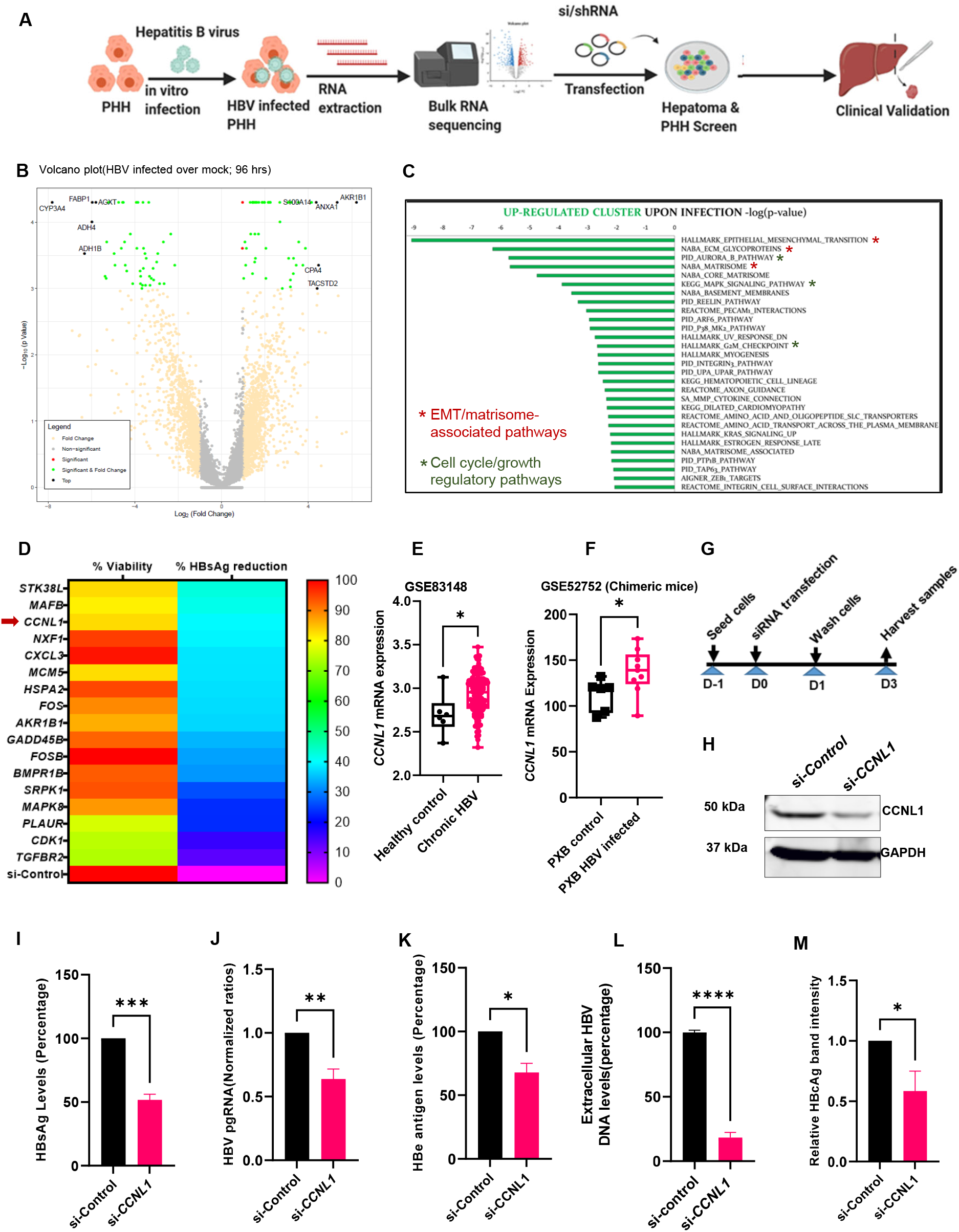
Identification of CCNL1 as a host susceptibility factor in HBV infection. **A**. Schematic methodology used to identify and validate the role of *CCNL1* during HBV infection. **B**. Volcano plot displays upregulated and downregulated genes upon HBV infection compared to mock infected PHH. **C**. Top upregulated pathways upon HBV infection. **D**. siRNA targeting each of the top 69 upregulated host factors were used in a HepAD38.7 RNAi screen to assess their role in HBV replication. Non-targeting control siRNA was included as a control. Hepatitis B surface antigen (HBsAg) ELISA was conducted on the cell culture supernatant, and cell viability was determined by CCK-8. Representative HBsAg reduction and cell viability are shown in the heat map. **E**. Expression of Cyclin L1 in healthy control and chronic HBV patient from micro-array dataset GSE83148. **F**. *CCNL1* expression in chronic HBV model of Phoenix Bio chimeric mice (GSE52572) is shown for the control and CHB. **G**. Experimental outline for the subsequent RNAi screen in HepAD38.7 cells. **H**. Knockdown of *CCNL1* is validated by Western Blot. **I-M**. Levels of HBsAg, HBV pgRNA, HBe antigen extracellular HBV DNA, and relative band intensity for the HBc antigen are shown. Statistical significance shown as p-value was analyzed by student t-test in GraphPad prism 8. Data are represented as mean ± SEM (n=3), ****p<0.0001, ***p<0.001, ** p<0.01, *p<0.05, ns; non-significant.

Although the gene expression data were consistent with some of the key pathways previously identified in the context of HBV infection, we also identified previously unsuspected players that have not been described. Therefore, we analyzed the function of the newly identified virally induced host genes in HBV replication and transcription. We designed an siRNA-based screen targeting the upregulated genes identified from WTS analysis. We used the HepAD38.7 cell line, which is easy to maintain in culture and contains HBV pgRNA under the control of a tetracycline-inducible on/off switch that can support HBV replication (Ladner *et al*, 1997; Hu *et al*, 2019; Ogura *et al*, 2014). The siRNA library consisted of three siRNAs per target gene, and a non-targeting siRNA was used as a negative control. From the HBsAg ELISA, we focused on candidate host factors whose knockdown resulted in a marked reduction in HBsAg levels without any appreciable effect on cell viability. This resulted in the identification of cyclin L1, whose RNAi-mediated loss-of-function resulted in a ∼40% reduction in HBsAg (**Fig. 1D**). Notably, the effect of CCNL1 on quantitative HBsAg levels was comparable with some of the previously described HBV host factors (also identified in our transcriptomic studies), including nuclear export transport factor (*NXF1*), which has been linked to the transport of pgRNA from the nucleus to the cytoplasm (Yang *et al*, 2014; Li *et al*, 2010), *CDK1* and cellular kinase R protein-specific kinase 1 (*SRPK1*), both of which have been linked to the phosphorylation of HBV core protein (Macovei *et al*, 2013; Ludgate *et al*, 2012; Daub *et al*, 2002; Heger-Stevic *et al*, 2018). To validate the clinical relevance of CCNL1 as a putative prognostic factor in CHB patients, we analyzed the expression of CCNL1 in liver tissues retrieved from the Gene Expression Omnibus database (**GSE83148, and GSE52752**). Remarkably, we observed consistent upregulation of *CCNL1* during chronic HBV infection (**Fig. 1E and F**). Furthermore, we observed enhanced expression of *CCNL1* in a cohort of local patients with unresolved CHB compared to those with resolved infection, as assessed by the loss of HBsAg (**Fig. S1B and B’**). Notably, analysis of the expression profiles in HBV e antigen-positive (HBeAg+) and-negative (HBeAg-) patients showed high CCNL1 expression in CHB patients irrespective of their HBeAg status (**Fig. S1C**). In contrast, patients who achieved a functional cure (loss of HBsAg) displayed significantly reduced expression of CCNL1 (**Fig. S1C**). Next, we evaluated the effects of cyclin L1 knockdown on HBV replication and gene expression in HepAD38.7 cells (**Fig. 1G**). The loss-of-function of Cyclin L1 was first validated by western blotting and rt-qPCR (**Fig. 1H and Fig. S2A)**, and, as shown in **Fig. 2B**, did not affect cell viability. Remarkably, *CCNL1* knockdown significantly reduced multiple viral parameters, including HBsAg, pgRNA, HBeAg, extracellular HBV DNA, and HBcAg ((**Fig. 1I-M, Fig. S2C)**. These results suggest that *CCNL1* may be an essential host factor that regulates HBV replication and transcription.

**Figure 2:**
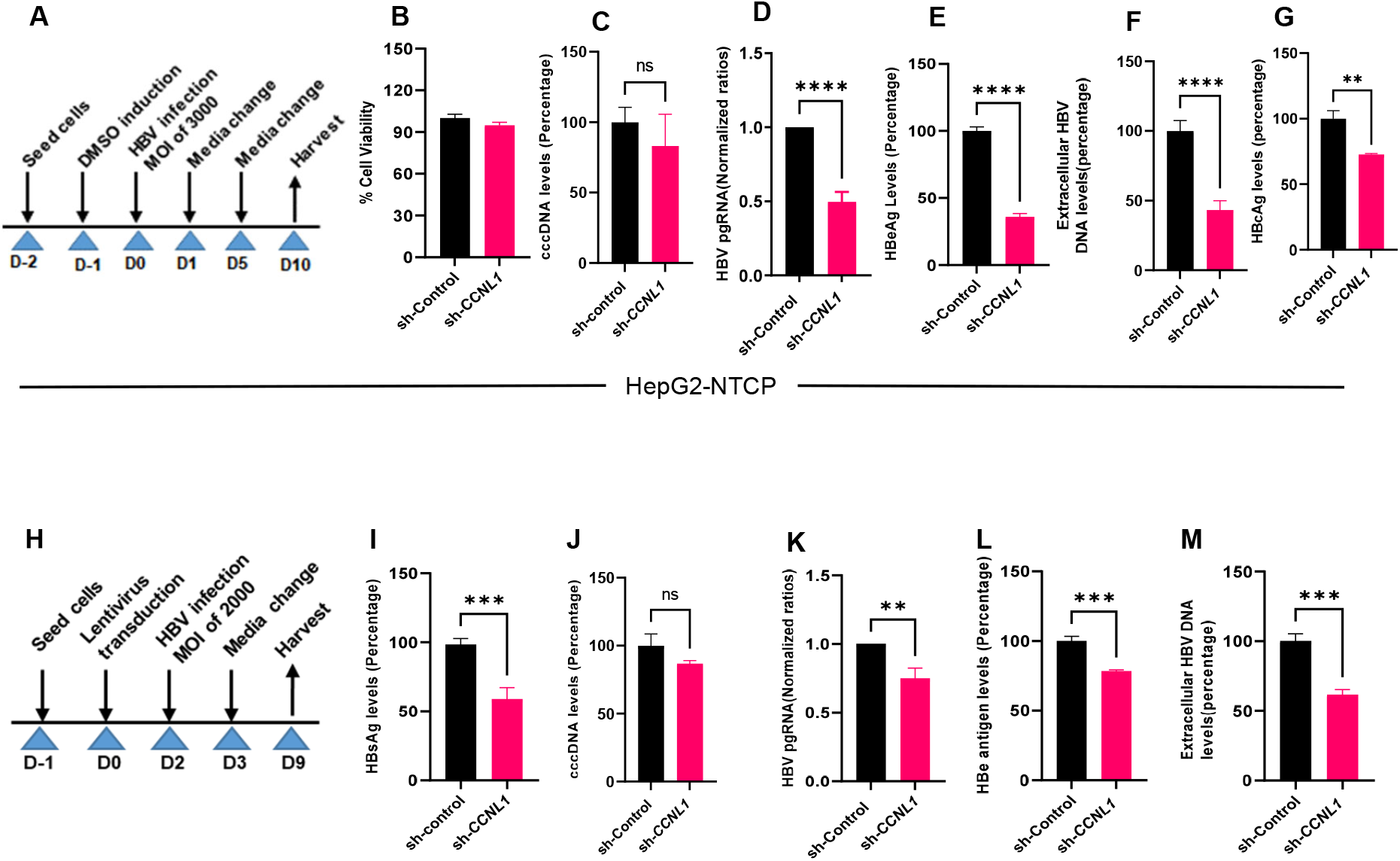
Functional validation of *CCNL1* in live infection models of HBV reveals its effect on distinct viral markers. **A**. Experimental outline in HepG2-NTCP is shown. **B**. Knockdown of *CCNL1* did not have any effect on the cell viability. **C**. No significant effect on the cccDNA is observed upon knockdown of Cyclin L1 in HepG2-NTCP. **D-F**. There was a significant reduction in pgRNA, HBeAg, and extracellular HBV DNA levels in *CCNL1* KD cells. **G**. HB core antigen (HBcAg) immunofluorescence analysis (IFA) shows the levels of infection in control and *CCNL1* knockdown cells. **G’**. Quantified HBcAg and normalized to the number of cells expressed as a percentage of the infection in control cells. **H**. Validation of the role of *CCNL1* during HBV replication in primary human hepatocytes. **I**. Knockdown of Cyclin L1 showed a remarkable reduction in surface antigen levels. **J**. Consistently, the knockdown of *CCNL1* did not show significant effect on cccDNA. **K-M**. Knockdown of *CCNL1* resulted in reduced levels of pgRNA, HBe antigen and extracellular HBV DNA. Statistical significance shown as p-value was analyzed by student t-test in GraphPad prism 8. Data are represented as mean ± SEM (n=3), ****p<0.0001, ***p<0.001, ** p<0.01, *p<0.05, ns; non-significant.

### Knockdown of cyclin L1 results in reduced HBV replication

To investigate the function of CCNL1 in a more physiological setting, we analyzed the effect of *CCNL1* knockdown on HBV replication and RNA synthesis in HepG2-NTCP (**Fig. 2A**) and PHH. CCNl1 knockdown in HepG2-NTCP cells was validated by WB (**Fig. S2D and quantified D’**). We also observed enhanced CCNL1 expression in infected HepG2-NTCP cells compared to that in controls (**Fig. S2E and E’**). Notably, the knockdown of cyclin L1 did not influence the viability of HepG2-NTCP cells (**Fig. 2B**), ruling out any non-specific or adverse effects. Interestingly, we did not observe any significant effect on HBV cccDNA levels upon *CCNL1* knockdown in HepG2-NTCP cells (**Fig. 2C**), suggesting that *CCNL1* likely functions downstream of cccDNA. Moreover, similar to the observations in HepAD38.7 cells, we found that loss-of-function of *CCNL1* in HepG2-NTCP cells also resulted in a remarkable reduction in HBV pgRNA, HBeAg, and extracellular HBV DNA (**Fig. 2D-F**). We also observed reduced levels of the hepatitis B core (HBcAg) protein in the knockdown cells compared to the controls (**Fig. 2G**). In contrast, the ectopic expression of *CCNL1* in HBV-infected HepG2-NTCP robustly enhanced the expression of HBV pgRNA, HBeAg, and extracellular HBV DNA (**Fig. S4A-F**), without any significant effect on cell viability or proliferation, suggesting a specific role in HBV control. To further bolster these findings, we assessed the role of CCNL1 in the HBV infection cycle in a live PHH infection model (**Fig. 2H-M**). The shRNA-mediated knockdown of Cyclin L1 was validated at the protein and mRNA levels (**Fig. S3A and 3B**). Similar to observations in other cell lines, knockdown of CCNL1 did not influence PHH viability (**Fig. S3C**). Remarkably, we observed significant inhibition of HBsAg (**Fig. 2I**), concomitant with a marked reduction in the levels of pgRNA (**Fig. 2K**), HBe antigen (**Fig. 2L**), and extracellular HBV DNA **(Fig. 2M**) in PHH transduced with CCNL1 shRNA. However, the effect of sh-CCNL1 on cccDNA levels remained insignificant, further corroborating the notion that *CCNL1* is likely to play a role in HBV control downstream of cccDNA formation. These results show that cyclin L1 is essential for HBV replication and gene expression without affecting the levels of cccDNA, possibly after the formation of cccDNA.

### CCNL1 modulates the production of HBV RNAs by regulating the activity of the total RNAPII

Given that *CCNL1* contains arginine-serine-rich dipeptide repeats, which are associated with pre-mRNA splicing (Dickinson *et al*, 2002), we evaluated the effect of *CCNL1* knockdown and overexpression on HBV splicing. We observed that the knockdown of *CCNL1* resulted in an overall decrease in the levels of HBV-spliced RNA variants, including singly spliced HBV RNA (Sp1) (**Fig. 3A; Fig. S5A and S5B**). In contrast, the ectopic expression of CCNL1 led to a global increase in spliced HBV RNA variants (**Fig. S5C and S6A**). Intriguingly, analysis of the singly spliced Sp1 RNA in the HBeAg+/-cohort of patients showed differential expression, with high levels of splice variants observed in HBeAg+, moderate levels in HBeAg-, and an almost complete absence in patients with HBsAg loss (**Fig. S5D**). Altogether, these results highlight the physiological relevance and association of CCNL1 expression with the clinical features of patients with CHB.

**Figure 3:**
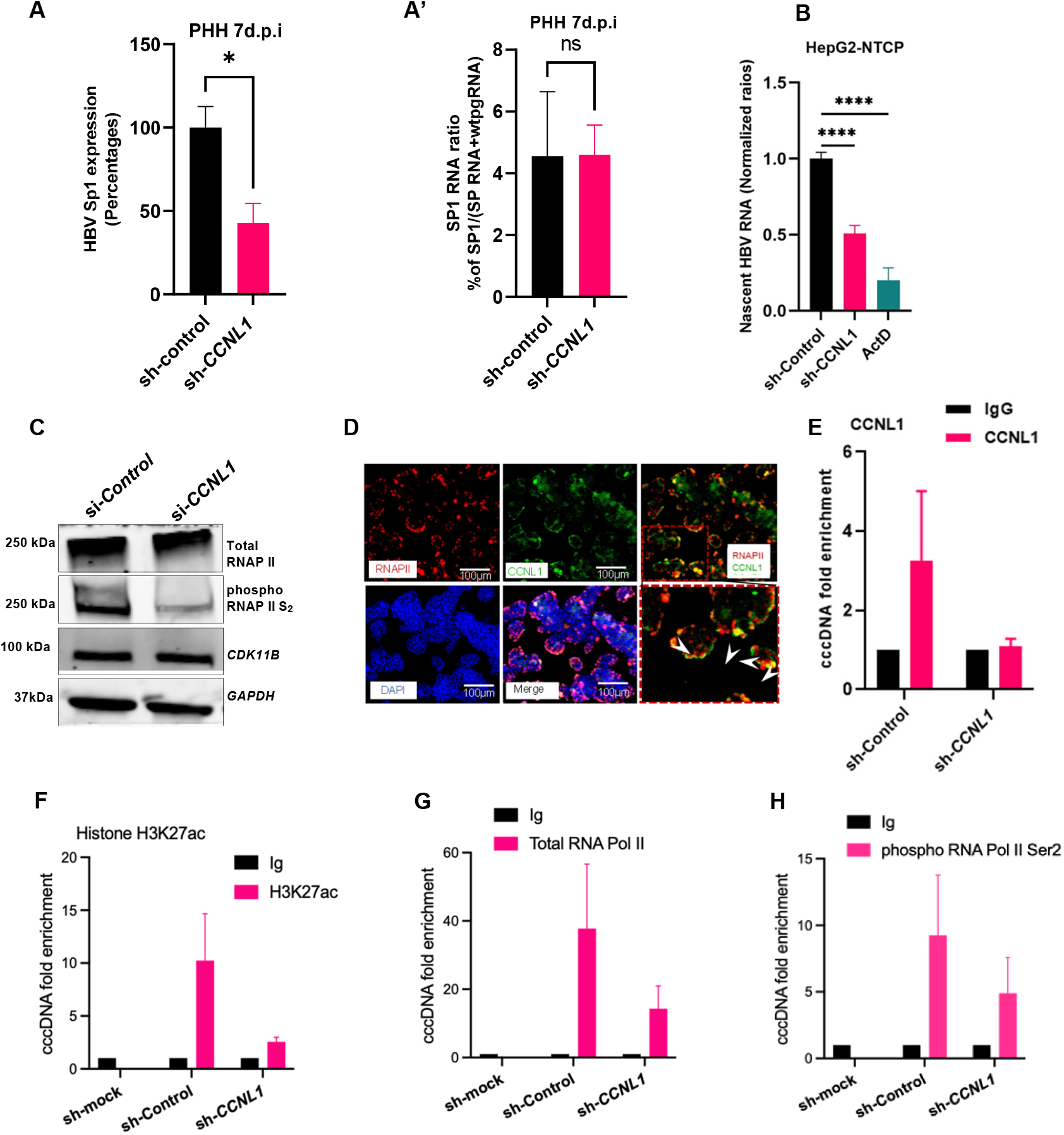
CCNL1 controls expression levels of HBV RNA by modulating cccDNA-dependent transcription. **A**. The HBV singly spliced RNA (Sp1) levels in the knockdown (shCCNL1) cells were expressed as a percentage of Sp1 levels in sh-control cells. **A’**. The splice ratio analysis wherein the levels of Sp1 in control and knockdown cells was calculated by dividing the expression of the Sp1 with the total HBV RNA+ Sp1 for the respective model in PHH. **B**. Knockdown of Cyclin L1 significantly reduced the production of nascent HBV RNA in HepG2-NTCP cells. Statistical significance shown as p-value was analyzed by student t-test for two groups in GraphPad prism 8. Data are represented as mean ± SEM (n=3), ****p<0.0001, *p<0.05, ns; non-significant. **C**. The effect of loss-of-function of *CCNL1* on total RNA polymerase II, phosphorylated S_2,_ and CDK11B is shown by Western Blot. **D**. Co-localization of Cyclin L1 and total RNAPII in HepG2-NTCP cells. The scale bar is 100μm. **E**. CCNL1 binds to cccDNA. **F-H**. The recruitment of RNA polymerase II and transcription mark on HBV cccDNA were detected by ChIP assay in HBV-infected HepG2-NTCP cells. cccDNA-ChIP was conducted in CCNL1 knockdown and control HepG2-NTCP cells infected with HBV **(F)**. The recruitment of H3K27ac on HBV cccDNA was significantly decreased after knockdown of CCNL1. As examined by ChIP-qPCR assay, CCNL1 knockdown remarkably reduced total RNA polymerase II recruitment on HBV cccDNA. RNAPII (**G**) and phosphorylated C-terminal domain serine 2 (**H**) associated-cccDNA were reduced in CCNL1-deficient cells. Shown data are representative of 3 independent experiments. Data are presented as the means ± SD.

However, given that our findings in HepAD38 and live infection models suggest a role for CCNL1 in controlling pgRNA levels, we wondered whether the alterations in the levels of splice variants resulted from alterations in the levels of pgRNA upon CCNL1 knockdown or overexpression. Indeed, normalization of the levels of splice variant Sp1 against pgRNA, both in the context of loss or gain of function of CCNL1, abrogated any of the observed absolute effects on splicing (**Fig. 3A’ and Fig. S6A’**). These results suggest that the impact of CCNL1 on the control of HBV splicing is likely to be indirect due to the regulation of pgRNA expression.

Next, we assessed whether Cyclin L1 influenced the production of HBV RNA. Indeed, siRNA- or shRNA-mediated knockdown of CCNL1 in HepAD38.7 (**Fig. S6B**) and HepG2-NTCP (**Fig. 3B**) cells resulted in a significant reduction in the levels of nascent HBV RNA. These results indicate that CCNL1 most likely plays a role in activating HBV RNA transcription. Previous studies have reported that *CCNL1* interacts with *CDK11B* to phosphorylate the C-terminal domain (CTD) of RNA polymerase II (RNAPII) (Maita & Nakagawa, 2020; Zhou & Fu, 2013). Indeed, siRNA-mediated knockdown of *CCNL1* resulted in markedly reduced phosphorylation of CTD-RNAPII at the Ser 2 (S_2_) residue (**Fig. 3C**), while overexpression resulted in increased S_2_ phosphorylation of CTD-RNAPII (**Fig. S6C**). Knockdown or ectopic expression of *CCNL1* did not significantly affect the expression of *CDK11B*, highlighting that the interaction did not affect the expression/stability of CDK11B (**Fig. 3C and Fig. S6C**). Additionally, immunofluorescence analysis revealed striking co-localization of *CCNL1* and total RNAPII (**Fig. 3D)**, further corroborating the putative function of *CCNL1* complexes in the regulation of RNAPII activity, which has been implicated in cccDNA-dependent viral transcription (Rall *et al*, 1983).

Next, we used a chromatin immunoprecipitation (ChIP) assay to assess whether CCNL1 can bind to cccDNA. Indeed, immunoprecipitation with an anti-CCNL1 antibody followed by PCR (ChIP-PCR) revealed that cyclin L1 can bind to HBV cccDNA, and the binding is diminished upon its knockdown (**Fig. 3E**). To assess the molecular consequences of CCNL1 knockdown on cccDNA-based transcription, we performed ChIP experiments using anti-H3K27ac, total RNAPII, phosphorylated RNAPII Ser2 and Ser5, and pan-acetylated histone H3ac. Interestingly, we observed a marked reduction in the binding of H3K27ac (a marker of active cccDNA-based transcription) to cccDNA upon CCNL1 knockdown (**Fig. 3F**). Additionally, consistent with the observed regulation of RNAPII phosphorylation, which was recently shown to regulate cccDNA-based viral transcription (Fan *et al*, 2022), our ChIP assays revealed that knockdown of CCNL1 also resulted in reduced binding of both total RNAPII and CTD-phosphorylated RNAPII to Ser2 (**Fig. 3G and H**) and Ser5 (**Fig. S6D**), to cccDNA. Furthermore, we observed a reduced interaction of pan-acetylated H3ac with cccDNA upon knockdown of CCNL1 (**Fig. S4E and F**). Altogether, these results suggest that CCNL1 may regulate HBV transcription by modulating the activity and recruitment of total RNAPII to cccDNA, and thereby, the subsequent downstream consequences of rewiring the epigenetic landscape of cccDNA by altering histone marks.

### Unbiased morphometric analysis of cellular processes altered by *CCNL1* knockdown in HepG2-NTCP cells

Thus far, our data suggest that CCNL1 could serve as a HBV host susceptibility factor regulating various aspects of the viral infection cycle, including viral transcription, and key cellular features vital to HBV replication. To further assess the phenotypic impact of CCNL1 knockdown on cellular features and cell-biological processes in an unbiased, holistic manner, we performed morphometric analysis with a Cell Painting assay using multiplexed fluorescent dyes (Bray *et al*, 2016) (**Fig. 4**). Principal component analysis (PCA) revealed two distinct cell clusters between sh-Control and shCCNL1 HBV-infected HepG2-NTCP cells (**Fig. S7**). Images from different channels were segmented into primary objects. Morphological, textural, and intensity features were extracted (**Fig. 4A**) using CellProfiler. The Pycytominer package was used to identify critical morphometric changes upon *CCNL1*. Notably, CCNL1 knockdown appeared to be associated with actin-and RNA-related processes, as well as intracellular transport, as suggested by the enrichment of Golgi-ER-related processes (**Fig. 4B**). To confirm these observations, we performed IFA to co-stain for the ER marker, protein disulfide isomerase (PDI), and CCNL1. Interestingly, we observed strong co-localization between the CCNL1 protein and PDI (**Fig. 4C**). These observations indicated possible perturbations in cellular processes upon CCNL1 knockdown. We speculated that some of the affected pathways may be essential for the HBV infection cycle. Further studies are required to validate these findings. Altogether, the data from our study suggest a working model whereby CCNL1 interacts with CDKs, such as CDK11B, to promote the phosphorylation of RNAPII. Consequently, phosphorylated RNAPII is recruited to cccDNA, where it regulates viral transcription (**Fig. 5**).

**Figure 4:**
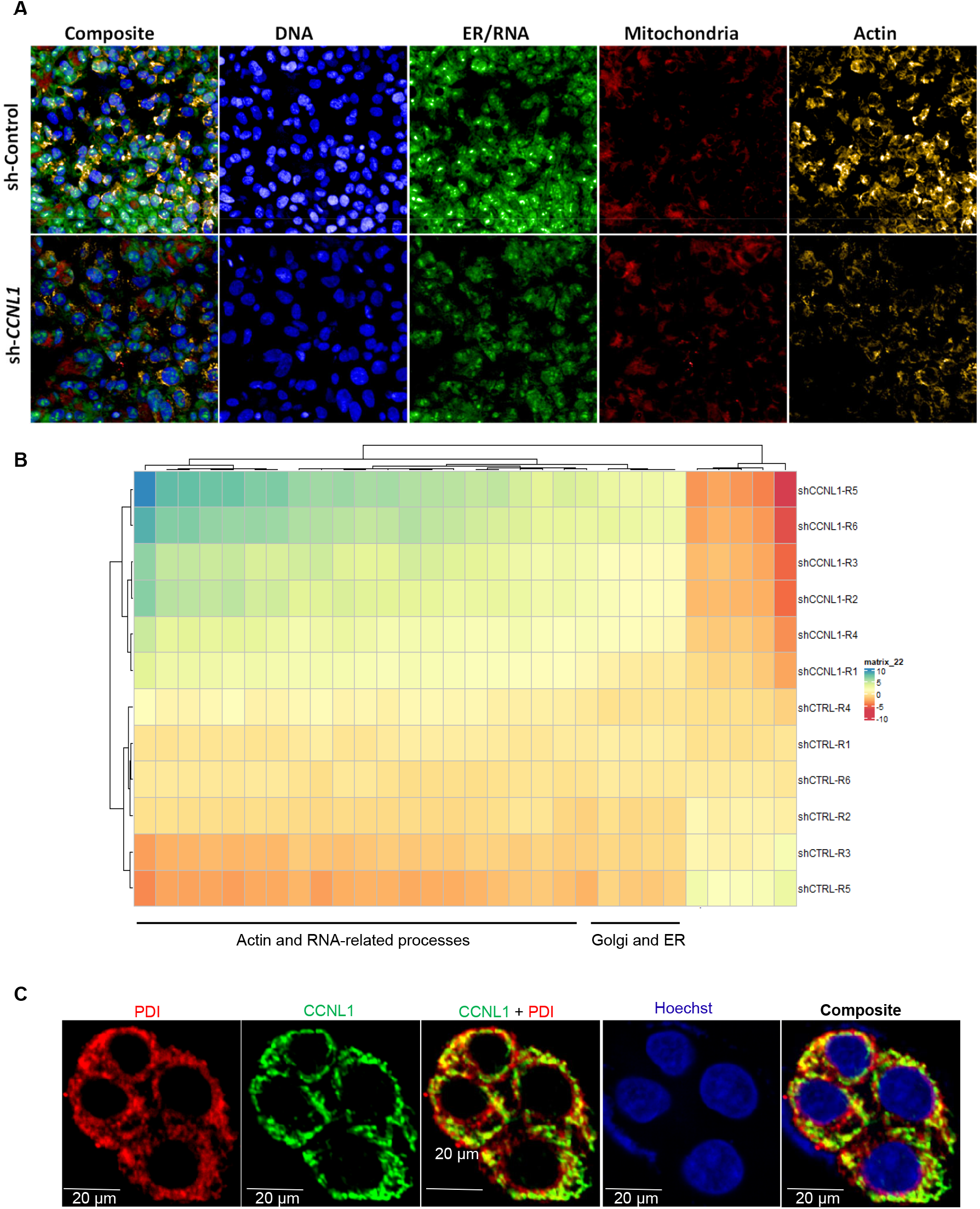
Morphometric analysis upon CCNL1 knockdown reveals phenotypic alterations in ER-related features that may associate with host pathways regulating the viral life cycle. **A**. Sample images from cell painting assay in HepG2-NTCP cells transfected with shControl or shCCNL1 and infected with HBV showing DNA, RNA/ER, mitochondria and Actin and composite are shown for the control and *CCNL1* knockdown cells. **B**. The top 30 differential cell features between control and knockdown cells are shown in the heatmap. **C**. Immunofluorescence analysis (IFA) image showing co-localization of ER marker, PDI, and *CCNL1* in HepG2-NTCP cells. The scale bar is 20μm.

**Figure 5:**
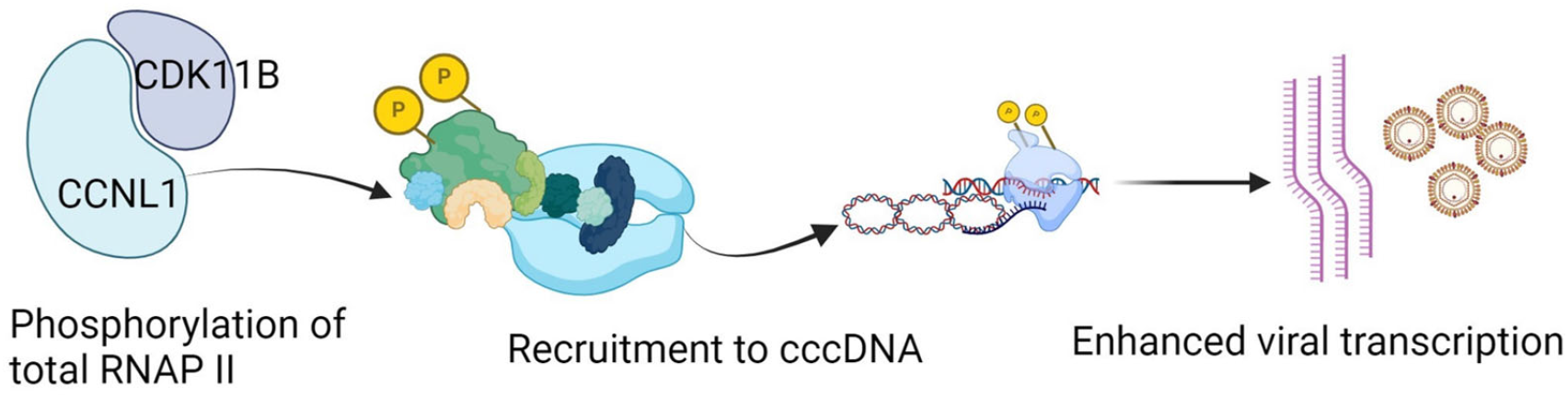
Working Model for HBV regulation by CCNL1. Here, we propose that cyclin L1, through interaction with CDK11B/A, phosphorylates total RNA polymerase II, which is then recruited to cccDNA. phosphorylated RNAP II controls cccDNA-dependent viral transcription, resulting in increased HBV gene expression.

## DISCUSSION

The hepatitis B virus, similar to other viruses, largely depends on host cell factors to establish an infection. Given that HBV has a high tropism to the human liver and mainly infects hepatocytes, the lack of optimal physiologically relevant, humanized models to study HBV-host cell interactions has remained a significant hurdle in the quest for identification of host-directed therapeutics (reviewed in (Hu *et al*, 2019)). Here, we performed RNA-seq analysis of HBV-infected PHH, which serves as a physiologically relevant model to identify putative host factors that are modulated and utilized by HBV to promote replication (**Fig. 1**). Numerous candidate host factors identified from this screen have been described in the literature as crucial during the HBV infection cycle, such as *SRPK1, CDK1*, and *NXF1* (Daub *et al*, 2002; Ludgate *et al*, 2012), thereby providing a crucial proof-of-concept for this approach.

Among the candidate host genes that were upregulated in PHH upon HBV infection, we focused on the downstream functional characterization of CCNL1. CCNL1 is known to bind to an atypical family member of cyclin-dependent kinases, CDK11. This study demonstrated that *CCNL1* may be a crucial host factor that supports efficient HBV replication and transcription. Specifically, siRNA-mediated knockdown of *CCNL1* results in a marked reduction in multiple viral parameters in heterologous cell models of HBV replication and live infection. Notably, we observed reduced extracellular HBV DNA and inhibition of HBV gene expression, including pgRNA, HBsAg, HBcAg, and HBeAg (**Fig. 1 and 2**). Given that *CCNL1* has been shown to interact with *CDK11B* to phosphorylate the CTD of RNAPII to modulate gene expression (Maita & Nakagawa, 2020; Zhou & Fu, 2013; Loyer & Trembley, 2020), we also investigated the effect of *CCNL1* on phosphorylation, and therefore on the activity of RNAPII. Indeed, the data from knockdown studies indicated that CCNL1 to control Ser 2 phosphorylation of CTD-RNAP II (**Fig. 3 and Fig.S6**). In this context, it is important to note that total and phospho-RNAPII have been shown to play crucial roles in the transcription of HBV from cccDNA (Fan *et al*, 2022).

Along the same lines of reasoning, we hypothesized that by influencing RNAPII activity, CCNL1 could modulate the transcription of HBV RNA (**Fig. 5**). Indeed, we found that CCNL1 knockdown resulted in reduced expression of the nascent HBV transcript, highlighting its importance in controlling viral transcription. Furthermore, ChIP-PCR assays showed that knockdown of *CCNL1* resulted in reduced H3K27 acetylation, a known marker of active cccDNA transcription (Pollicino *et al*, 2006; Yang *et al*, 2020) as well as the binding of total RNAPII and RNAPII S_2_ to cccDNA (Fan *et al*, 2022). Previous studies have shown that CCNL1 forms a complex with SC35/SRSF2, CDK11, *and* CDK12 to regulate pre-mRNA splicing (Chen *et al*, 2006; Dickinson *et al*, 2002; Loyer & Trembley, 2020). Interestingly, spliced HBV RNA variants have been identified in the serum and liver tissues of patients with chronic HBV (Soussan *et al*, 2000; Sommer & Heise, 2008; Candotti & Allain, 2016; Lee *et al*, 2008). These spliced variants and products have been linked to play a role as immune decoys, associated with liver damage, fibrosis, and the development of HCC (Duriez *et al*, 2017; Soussan *et al*, 2003; Preiss *et al*, 2008; Pol *et al*, 2015; Wu *et al*, 2018; Chen *et al*, 2015; Bayliss *et al*, 2013). Thus, we analyzed the HBV splice variants in our models and found that knockdown of CCNL1 resulted in a global decrease in HBV variants in hepatoma cells. However, given our observation that CCNL1 controls HBV transcription, normalization of the levels of spliced HBV RNA against total HBV RNA or pgRNA failed to reveal any appreciable difference in the levels of splicing upon loss-of function of CCNL1. Altogether, our data suggest that while CCNL1 may have an indirect role in modulating the levels of HBV RNA splice variants by regulating the overall transcription of HBV, it is less likely to play a direct role in pre-mRNA splicing of nascent HBV RNA (**Fig. 3**). In addition, knockdown of CCNL1 resulted in a decreased association of total and Ser2-phosphor RNAPII with cccDNA, further highlighting its role in regulating cccDNA transcription. These observations corroborate recent reports suggesting that the regulation of total RNAPII phosphorylation affects cccDNA-based viral transcription (Fan *et al*, 2022). Our studies showed that CCNL1 co-localizes with total RNAPII and likely recruits it to the cccDNA mini-chromosome in a phosphorylation-dependent manner (**Fig. 3**).

Unbiased morphometric profiling of CCNL1 perturbation in HepG2-NTCP cells (**Fig. 4**) revealed that *CCNL1* might be involved in regulating diverse cell features critical for HBV replication, such as actin, the endoplasmic reticulum, and the Golgi apparatus. Notably, HBV has been reported to modulate host actin polymerization, thereby affecting the morphology of hepatoma cells (Kim *et al*, 2021; Kong *et al*, 2017). The HBx protein was shown to specifically interact with calmodulin (CaM), resulting in the displacement of HSP90, activating the cofilin pathway, thereby enhancing the polymerization of actin, and has been linked to HBV-mediated HCC metastasis (Lara-Pezzi, 2001; Kong *et al*, 2017; Kim *et al*, 2021). The core protein was also reported to interact with the C-terminal domain of actin-binding proteins (Huang *et al*, 2000). These observations highlight the diverse putative functions of CCNL1 in modulating key HBV-dependent cell biological phenotypes that may be relevant for CHB-driven chronic liver disease and cancer. We also observed colocalization of CCNL1 with the ER marker PDI, whose member ERp57 was recently shown to play a vital role in regulating HBV membrane fusion and infection (Pérez-Vargas *et al*, 2021). The exact role of *CCNL1* in host cell features that facilitate the HBV life cycle remains to be elucidated in future studies.

Finally, our study revealed a robust correlation between the enhanced expression of *CCNL1* and chronic HBV infection in clinical samples (**Fig. S1B and B’**). Specifically, patients exhibiting a functional cure, as assessed by loss of HBsAg, displayed much lower CCNL1 expression, thereby corroborating its clinical relevance as a prognostic biomarker for CHB patients in the clinic. Overall, this study not only contributes to a better understanding of HBV infection biology, but also offers putative druggable targets for the development of novel, host-directed therapies against HBV.

## MATERIALS AND METHODS

### Cell culture and treatments

The HepG2-NTCP cells used in this study were a kind gift from Assoc Prof Tan Yee Joo, NUS, and have been reported to be susceptible to HBV infection (Iwamoto *et al*, 2014). HepG2-NTCP cells were maintained as described earlier (Makokha *et al*, 2019). HepAD38.7-Tet cells used in this study were cultured as previously described (Ogura *et al*, 2014). HEK293Tcells were maintained in antibiotic-free DMEM/F12+Glutamax (Gibco) supplemented with 10% FBS. PHH were purchased from Lonza and maintained according to the manufacturer’s instructions.

Lentiviral transduction of the cells was carried out in the presence of 5 μg/ml polybrene (sc-134220, Santa Cruz), followed by selection of positive clones in the presence of 10 μg/ml puromycin (ant-pr-1, InvivoGen). All plasmid and siRNA transfections were performed using Lipofectamine 3000 (Invitrogen) according to the manufacturer’s protocol.

### Production of HBV stock virus for infection

The virus used for infection in this study was produced as described earlier (Lim *et al*, 2022). Briefly, cell culture supernatant was harvested from tet-off-HepAD38.7 cells and filtered through a 0.22 µm polyethersulfone (PES) membrane filter. This was followed by concentration using a heparin column and elution with high salt, followed by dialysis in phosphate-buffered saline overnight.

### Generation of stable cell lines

Flag-tagged plasmids expressing *CCNL1*, and empty vector open reading frames (ORFs) were sourced from GenScript. Transfection was performed using Lipofectamine 3000 (Invitrogen) with 9 μg of plasmid in 10 cm cell culture dishes. Cells expressing the plasmids were selected using DMEM supplemented with 10% FCS and 10 μg ml^−1^ blasticidin (Santa Cruz Biotechnology).

### Plasmids, antibodies, oligos, siRNA, and shRNA

The oligonucleotides used in this study were designed using Primer Plus and synthesized by Integrated DNA Technologies, Inc. (IDT). Validated siRNAs targeting the three regions of the upregulated genes were ordered from Silencer Select (Applied Biosystems). Control shRNAs and shRNAs targeting *CCNL1* were obtained from Thermo Fisher Scientific. Access to these libraries was provided by high-throughput phenomics (EDDC). The following packaging vectors were used in this study: pMD2.G (Addgene plasmid #12259), pRSV-Rev, and pMDLg/pRRE, all gifts from Didier Trono (Dull *et al*, 1998). A comprehensive list of the antibodies used in this study is shown in Table S1.

### RNA-seq of infected primary human hepatocytes

Fresh primary human hepatocytes (Invitrocue/Yecuris) from FRG KO humanized mice were infected with HBV from a cell culture (HepAD38.7 cells) at an MOI of 3000. Cell lysates were collected 96 h post-infection. RNA isolation was performed using an RNeasy kit (Qiagen, Hilden, Germany). Quality was assessed using a Bioanalyzer, the library was prepared using SureSelect XT Human All Exon V6, and sequencing was performed using Novaseq 6000 (Theragen). Differentially expressed genes between mock-infected and HBV-infected samples were identified by DEGseq analysis in R. Gene set enrichment and pathway analyses were performed using DAVID software to determine the top upregulated and downregulated pathways, and the top genes were identified for further analysis.

### Immunofluorescence staining and confocal microscopy

The cells were seeded in 96 well plates (PerkinElmer Ultra plates) and treated as previously described. Cells were fixed in 4% paraformaldehyde at room temperature for 10 min, followed by permeabilization with 0.1% Triton-X for 5 min at room temperature. Fixed cells were blocked with 3% bovine serum albumin solution for 5 min at room temperature and incubated with primary antibodies overnight at 4 °C. The cells were washed thrice with 1x PBS to remove excess primary antibodies. The secondary antibody incubation was carried out for 1 h at room temperature before the cells were washed and counterstained with DAPI (Sigma Aldrich). The plates were imaged using the Operetta CLS confocal microscope (PerkinElmer). Infected cells were quantified using the Columbus Data Storage and Analysis System (PerkinElmer). The data were normalized to the number of cells per well and the control wells were equated to 100%.

### HBV infection of HepG2-NTCP and PHH

HepG2-NTCP and PHH cells were seeded in type I collagen (R-011-K, Gibco)-coated plates. HepG2-NTCP cells were infected at an MOI of 3000 in the presence of 4% PEG8000 and 1% DMSO at 37°C in a 5% CO_2_ incubator, as described previously (Dull *et al*, 1998). Primary human hepatocytes were infected at an MOI of 2000 in the presence of 4% PEG8000 and 1% DMSO, as described previously (Xiang *et al*, 2019). The samples, cell lysate for RNA and DNA, as well as cell culture supernatants, were harvested at the indicated time points, as shown in the figure legends.

### RNAi screening in HepAD38.7 cells

HepAD38.7 cells were maintained as previously described. A pool of three siRNAs per gene was spotted into 384 well plates at a final concentration of 20nM. Gene silencing was performed by reverse transfection using the Lipofectamine 3000 reagent protocol (0.1 μL Lipofectamine 3000 per well in Opti-MEM medium) (ThermoScientific). The cells were washed, and the media changed one day post-transfection. Transfected cells were incubated at 37°C in a 5% CO2 incubator for 72 h. For reproducibility, the screen was independently repeated three times. Four technical replicates were included for each plate, and two biological replicates were included for each screen.

### Quantification of HBsAg, HBeAg, and extracellular HBV DNA

The cell culture supernatant was collected at the end of the experiment. For HBsAg ELISA, 384 well polystyrene plate Maxisorp Cat# P6366-1CS (Sigma-Aldrich) was coated with 6 μg/ml mouse monoclonal anti-HBsAg (ab20758 at 6 μg/ml) in 0.05M carbonate-bicarbonate buffer pH 9.6 (Cat# C3041-100, Sigma Aldrich) and incubated overnight at 4°C. The plates were thawed and washed three times with wash buffer (0.05% Tween 20 in 1x PBS) before blocking in assay diluent (10% FBS in 1x PBS), followed by incubation for 2 h at room temperature. The plates were washed three times before adding 25 μl of sample and standard (HBsAg (Adw) full-length protein, ab91276) and incubated for 2 h at room temperature. Plates were washed five times before adding rabbit polyclonal anti-biotin (ab68520 1:1000 dilution) and streptavidin-HRP (cat#554066, BD 1:1000) followed by 1-hour incubation at room temperature. The plates were washed 7 times to remove any excess working detector, followed by adding TMB substrate solution (Pierce, ThermoScientific) and incubated at room temperature for 30 minutes. The reaction was stopped by adding 1N HCl stop solution before reading the absorbance at 450 nm using a Tecan 100 spectrophotometer. HBsAg levels are expressed as a percentage of the control set at 100%. HBeAg was quantified following a one-step sandwich CLIA protocol, according to the manufacturer’s instructions (Autobio Diagnostics). Wild-type and defective HBV DNA was amplified from DNA extracted from 200μL of cell culture supernatant from HBV-infected HepG2-NTCP, PHH, and HepAD38.7, as previously described (Preiss *et al*, 2008; Bayliss *et al*, 2013).

### Quantitative Real-Time PCR (qPCR)

RNA was extracted from the cell lysates according to the Total RNeasy kit protocol (Qiagen). cDNA was synthesized from 1μg of extracted RNA using Invitrogen SuperScript IV VILO Master Mix (Cat. 11766500, Life Technologies). HBV pre-genomic RNA and gene mRNA levels for knockdown were determined using the SYBR Fast-based qPCR master mix kit (KAPPA BIOSYSTEMS) in the Quant studio 7 PCR system (Applied Biosystems). The pgRNA primers used were as follows: forward, CGTTTTTGCCTTCTGACTTCTTTC and reverse, ACAGAGCTGAGGCGGTGTCTA (Wang *et al*, 2015). qPCR data were analyzed using the Livack method as described previously (Livak & Schmittgen, 2001). All primers used in this study for quantitative polymerase chain reaction (qPCR) were purchased from Integrated DNA Technologies.

### Quantification of cccDNA using TaqMan probe-based qPCR

HBV cccDNA was quantified as described earlier (Ciechanover *et al*, 1982; Huang *et al*, 2018). Briefly, DNA was extracted at the end of the experiment according to the DNA mini kit (Qiagen) protocol with proteinase K digestion for 1 h at 37°C. 1μg of the extracted intracellular DNA was digested with T5 exonuclease (New England Biolabs) to remove the relaxed circular and single-stranded HBV DNA according to the manufacturer’s protocol (Qu *et al*, 2018). Briefly, digestion was performed at 37°C in the presence of 10 units of T5 exonuclease and NEBuffer 4 for 30 min. The reaction was stopped by adding 11 mM EDTA. Quantification of cccDNA was performed using the specific primer Fwd-primer: 5’-GGGGCGCACCTCTCTTTA-3’ ccDNA Rev-primer: 5’-AGGCACAGCTTGGAGGC-3’ TaqMan probe: 5′-FAM-TCACCTCTGCCTAATCATCTC-TAMRA-3’ (Huang *et al*, 2018). To enhance the specificity of the PCR, 100 nmol/l forward and 400 nmol/l reverse primers were used. Denaturation was conducted at 95°C for 10 min, followed by 40 cycles of step 1 (95°C for 1 s) and step 2 (65°C for 1 min), as previously reported by Allweiss et al. (Allweiss *et al*, 2018).

### Western blot analysis

Protein samples were separated by SDS–PAGE, and then transferred to PVDF membrane, transferred to PVDF membranes, blocked with Intercept^™^ (TBS) buffer (Bio-Rad, P/N: 927-70001) or 5% skim milk for 1 hour at room temperature and incubated with primary antibodies at 4°C overnight. Following incubation with the appropriate secondary antibodies (LI-COR IRDye® 800CW anti-rabbit 92632211 (LiCor, 1:5000) or 680CW anti-mouse 9268070 (LiCor, 1:5000)), the blots were visualized using the ChemiDoc XRS+system (Bio-Rad).

### ChIP assays

Chromatin immunoprecipitation (ChIP) was performed using the MAGnify Chromatin Immunoprecipitation System (# 492024; Applied Biosystems). The cells were fixed with 1% paraformaldehyde (PFA) for 10 min at room temperature. Fixation was stopped by adding a 1.25M glycine solution for 5 min at room temperature. Cells were subsequently washed in chilled PBS (Eurobio), resuspended in ChIP lysis buffer supplemented with protease inhibitors at 1 million cells/50 mL, and incubated for 5 min on ice. The chromatin solution was then sonicated for 15 cycles of 30 s ON and 30 s OFF using a Picoruptor Sonicator (Diagenode) to generate 200- to 500-bp DNA fragments. The debris was pelleted by centrifugation at 20,000 × g and 4°C for 5 min. The sheared chromatin was diluted to a concentration of 200,000 cells/100 mL and incubated overnight at 4°C with the previously coupled antibody-Dynabeads protein A/G under end-over-end rotation. Ten microliters of mouse anti-RNAPII, RNAPII Ser2P, RNAPII Ser5P, and rabbit anti-H3K27ac antibodies (Active Motif) were used (Table **S1**). Immunoprecipitation with non-specific immunoglobulins (Applied Biosystems) was included in each experiment as a negative control. After washing and reverse crosslinking, immunoprecipitated chromatin was purified using DNA purification magnetic beads (Applied Biosystems), according to the manufacturer’s protocol. Purified DNA was analyzed by real-time quantitative PCR using specific primers and fluorescent probes for HBV cccDNA (primers CCGTGTGCACTTCGCTTCA / GCACAGCTTGGAGGCTTGA; probes CATGGAGACCACCGTGAACGCCC).

### Viability/cytotoxicity assays

Cell viability and cytotoxicity were evaluated using the CellTiter-Glo Luminescent assay (Promega) and cell-count kit-8 (Dojindo, Japan), respectively, following the manufacturer’s instructions.

### Cell painting Assay

Stable knockdown and control HepG2-NTCP cells were seeded in 384 well plates and infected with HBV as described previously. The detailed staining protocol is described in the supplementary file. The cells were fixed and stained with cell painting dyes followed by imaging with a Perkin Elmer Operetta CLS, and image segmentation and feature extraction were performed using CellProfiler pipelines adapted from Bray et al. (26). The dyes that were used are listed in Table S2. Cell painting analysis pipelines using pycytominer and R scripts were developed at the High Throughput Phenomics Lab at EDDC.

### Statistical analysis

Data analysis was performed using GraphPad Prism 8 using different tests, as indicated in the figure legends. Network analysis of the upregulated genes was performed using the STRING database, as previously described (Szklarczyk *et al*, 2017). Differentially expressed genes from the RNA-seq data were analyzed in R studio using the DEGseq package, as reported previously (Wang *et al*, 2009).

## Author contributions

Conceptualization, RD, JPB, COO, BCN, GP, ML and SGL.; Methodology, RD, JPB, COO, BCN and MLP; Investigation, COO, BCN, MLP, PPKA, NG, JTS, GYL and TTM.; Formal Analysis, BCN, COO, NS and TTM.; Writing – Original Draft, COO, BCN and RD.; Writing-Review and Editing, RD, JPB, COO, and BCN.; Funding Acquisition, RD and SGL.; Resources, SGL and RD.; Supervision, RD, JPB, SGL, ML and BCN.

## Acknowledgments

The authors thank Assoc Prof. Tan Yee Joo (Institute of Molecular and Cell Biology, Agency for Science Technology and Research, Singapore 138673, Singapore) for sharing reagents and viruses used for infection in this study. We thank Prof. David Durantel and Dr. Anna Salvetti of CIRI INSERM, Lyon, for sharing their protocols and for their support, insight, and guidance. We thank Koichi Watashi and Takaji Wakita (National Institute of Infectious Diseases, Japan) for providing reagents. We thank Prof. Adam Zlotnick for sharing the anti-HBc antibodies. We thank Ms. Ming Jie Lim for her assistance with the siRNA screening. We thank Assoc Prof. Tan Yee Joo for producing the virus stocks used in the infection experiments. We thank Weiyong Chua for providing the administrative support. Lastly, we thank all patients whose samples were used for validation in this study.

