## Supplementary methods for "Identification of Cyclin L1 as a host factor regulating Hepatitis B Virus replication"

**Fig. S1 Analysis of HBV mediated transcriptomic changes**

**A.** Down-regulated pathways upon HBV infection in PHH at 96hrs compared to the mock control. RNA or protien was extracted from liver tissue from CHB patients and patient who have achieved surface antigen loss (HBsAg negative) and HBV pgRNA quantified by RT-qPCR and the expression normalized to beta actin. **D.** Protein was extracted from the liver tissues and 30μg of RNA analyzed in Western Blot to show the levels of *CCNL1* in HBsAg negative and CHB samples. **B’.** Quantification of band intensity for *CCNL1* normalized to the beta actin levels for the CHB and HBsAg negative patients. **C.** Levels of *CCNL1* mRNA in HBsAg negative, HBeAg- and HBeAg+ samples quantified by RT-qPCR and normalized to the levels of beta actin.

**Fig. S2 Evaluating the role of *CCNL1* during HBV infection**

**A.** Quantification of the CCNL1 band intensity and normalized to GAPDH housekeeping gene. **B.** siRNA knockdown of Cyclin L1 did not show toxicity in HepAD38.7 cells. **C.** Representative Western Blot image showing the levels of the HBV core protein. **D.** Validation of the *CCNL1* knockdown by shRNA. **D’.** Quantification of CCNL1 band intensity and normalized to the GAPDH levels and expressed as a percentage to the control cells. **E**. IFA staining of CCNL1 in HBV-infected cells compared to control cells. **E’**. Quantification of the CCNL1 expression intensity in infected vs control cells. Statistical significance shown as p-value was analyzed by student t- test in GraphPad prism 8. Data are represented as mean ± SEM (n=3), ****p<0.0001, ***p<0.001, ** p<0.01, *p<0.05, ns; non-significant.

**Fig. S3 Functional validation of Cyclin L1 in PHH**

**A.** Validation of Cyclin L1 knockdown in PHH by Western Blot. **B.** Levels of *CCNL1* at mRNA level by RT-qPCR. **C.** Bright field images of sh-*CCNL1* and sh-control transfected PHH at day 9 (6dpi) and 10 (7dpi) in culture. The scale bar is 50μm. Statistical significance shown as p-value was analyzed by student t- test in GraphPad prism 8. Data are represented as mean ± SEM (n=3), ****p<0.0001, ***p<0.001, ** p<0.01, *p<0.05, ns; non-significant.

**Fig. S4 Ectopic expression of *CCNL1* in HepG2-NTCP cells enhances HBV gene expression**

**A.** Cell viability for the control and cells overexpressing Cyclin L1. **B.** Expression of *CCNL1* at mRNA levels as quantified by RT-qPCR. C. Validation of ectopic expression of CCNL1 by Western Blot. **C’.** Quantified CCNL1 band intensity for the control and overexpression cells and expressed as a percentage to the control. **D.** Levels of HBV pgRNA in control and HepG2-NTCP cells overexpressing *CCNL1* are shown. **E.** Levels of HBe antigen quantified by CLIA are shown. **F**. HBV extracellular DNA levels are shown as quantified by qPCR. Statistical significance shown as p-value was analyzed by student t- test for two groups and one-way ANOVA for more than two groups in GraphPad prism 8. Data are represented as mean ± SEM (n=3), ****p<0.0001, ***p<0.001, ** p<0.01, *p<0.05, ns; non-significant.

**Fig. S5: Evaluating the role of Cyclin L1 on HBV RNA splicing**

**A-C.** Levels of HBV spliced RNA variants in HepG2-NTCP and HepAD38.7 in both knockdown and overexpression models. **D.** Agarose gel displaying products of RT-qPCR of HBV Sp1 in the three categories of patients.

**Fig. S6: Cyclin L1 interacts with HBV RNA**

Evaluation of RNA stability in Huh7 cells. Huh7 cells expressing empty vector or *CCNL1* ORF were transfected with Tet-HBV and cells harvested 72 hours later (timepoint 0), Tetracycline was added, and samples were harvested 6- or 8-hours post-addition of tetracycline. The Levels of the HBV RNA was expressed as a percentage to the amount captured at timepoint 0. **A.** Levels of remaining HBV RNA 8 hours post-addition of Tetracycline. **B and C.** RNA immunoprecipitation (RIP) using anti-*CCNL1* antibody and IgG control were used for pulldown and the levels of S, X and core RNA quantified by RT-qPCR from Huh7-transfected and HepG2-NTCP models respectively. Data are represented as mean ± SEM (n=3), ****p<0.0001, ***p<0.001, ** p<0.01, *p<0.05, ns; non-significant.

**Fig. S7 Morphometric analysis of HepG2-NTCP cells after Cell Painting and hierarchical clustering display two different clusters of sh-*CCNL1* and sh-control cells**

PCA showing distinct clusters of sh-control and sh-*CCNL1* knockdown cells from cell painting analysis in HepG2-NTCP infected with HBV.

**Methods: Supplementary**

**Nascent RNA Assay**

Actinomycin D (Invitrogen, A7592) was added to the culture media of the wells at a final concentration of 0,79 μM (1/1000, photosensitive) and incubated for 1-hour. After 1 hour at 37°C, 5-bromouridine (BU, SIGMA 850187) was added to culture media at a final concentration of 2 mM for 2h at 37°C. After the 2-hours incubation, the media was aspirated, and the cells washed 1X with PBS and thereafter lysed the cells with 300 μl of RA1 + DTT. RNA was extracted according to Qiagen RNA extraction kit protocol. In the meantime, 25 μl of Dynabeads goat anti-mouse IgG (Life Technologies, 1033) per sample was washed with PBS-BSA 0.1% supplemented with SUPERaseIn (Invitrogen, AM2696). After the washing, 1 μg of anti-BrdU antibody (BD Pharmingen 555627) was added and further incubated for 1 hour at room temperature on a rotary agitator. 1 μg of total RNAs was incubated for 10 minutes at 80°C and immediately put tubes on ice thereafter. The total RNAs were precleared with 5 μl of Dynabeads in a final volume of 400 μl of PBS-BSA 0.1% + SUPERaseIn and incubate 1 hour at 4°C on a rotary agitator. The beads were collected on a magnet and the supernatant incubated with Dynabeads conjugated with BrdU antibody overnight at 4°C on a rotary agitator. After overnight incubation, the beads were washed 3X 5 minutes with PBS-BSA 0.1% + SUPERaseIn. The nascent RNAs were eluted by adding 100 μl of water + SUPERaseIn and incubated for 10 min at 95°C. The nascent RNAs were purified and concentrated following the RNA clean and concentrator kit (Zymo Research, ZR1017) instructions and the RNAs eluted in 25 μl final volume. The RNAs were reverse transcribed using Superscript VILO cDNA synthesis kit (Cat# 11754050, Invitrogen) and qPCR performed as described earlier using SYBR, pgRNA primers and RPLPO as a control.

**Cell Painting Protocol**

To evaluate the effect of *CCNL1* knockdown on HepG2-NTCP cell morphology, we carried out a cell painting assay as described earlier ^1^. Step by step protocol is described below

1. Seed cells into 384-well plate in growth media (50 µL cell suspension/well) at a density of 600 cells/ well
2. Incubate overnight at 37°C, 5 % CO_2_ to allow overnight recovery and growth of plated cells
3. Incubate the plates at 37°C, 5% CO_2_ for 48 hours
4. Prepare MitoTracker staining solution from 1 mM stock:
   a. Add 3 µL of 1 mM stock per 1 mL of media
5. Do NOT remove media from wells! Add 10 µL of diluted MitoTracker staining solution and spin at 500g for 1 minute
6. Incubate in the dark for 30 minutes at 37C, 5% CO_2_ (in incubator)
7. Add 20 µL of 16% PFA to achieve final concentration of 4% PFA and spin at 500g for 1 minute
8. Incubate for 20 minutes at RT (in drawer)
9. Remove supernatant and wash 2x with 1X HBSS, 70 µL/well (
10. Prepare permeabilization & staining solution (1x 384-well plate: 10ml):
    a. 0.1% Triton X-100, 1% BSA (w/v) in 1X HBSS (10 mL)
       - Since BSA may contain contaminants, prepare filtered 3% BSA in HBSS solution
    b. Phalloidin (12.5 µL)
    c. Concanavalin A (10 µL)
    d. Hoechst (1 µL)
    e. WGA (15 µL)
    f. SYTO14 (12 µL)
11. Add 20 µL of permeabilization & staining solution, incubate for 30 min at RT
12. Discard solution and wash 2x with 1X HBSS, 70 µL/well
13. Fill the wells with 60 µL of 1X HBSS or 0.05% Sodium azide (for long term storage)
14. Seal plates tightly with adhesive seals

**Analysis of HBV spliced RNA variants**

The fifteen known HBV splice variants were analyzed as described recently by Chabrolles and colleagues ^2^. Quantitative qPCR was performed to detect HBV variant RNA including the unspliced RNA and the data normalized to the GAPDH and beta-actin genes in both knockdown and ectopic expression of *CCNL1*. The primers that detect all 15 spliced RNA (sv1 to sv15) and intron1, 2, and 2b were designed by Sommer and colleagues ^3^. To further analyze splicing events upon knockdown or ectopic expression of *CCNL1,* the singly spliced HBV RNA (sp1) primers (SP1, 5′‐TGCCCCTATCTTATCAACAC‐3′ (nt 2311–2330, sense) and SP2, 5′‐CAAATGGCACTAGTAAAC‐3′ (nt 695–678, antisense)) were used to assess spicing as described earlier by Sommer and colleagues ^3^. These primers were also used to assess the levels of HBV spliced variant RNA in patient samples.
