## Supplementary figures for "Identification of Cyclin L1 as a host factor regulating Hepatitis B Virus replication"

**Fig. S1 Analysis of HBV mediated transcriptomic changes and CCNL1 expression CHB and infected cells**

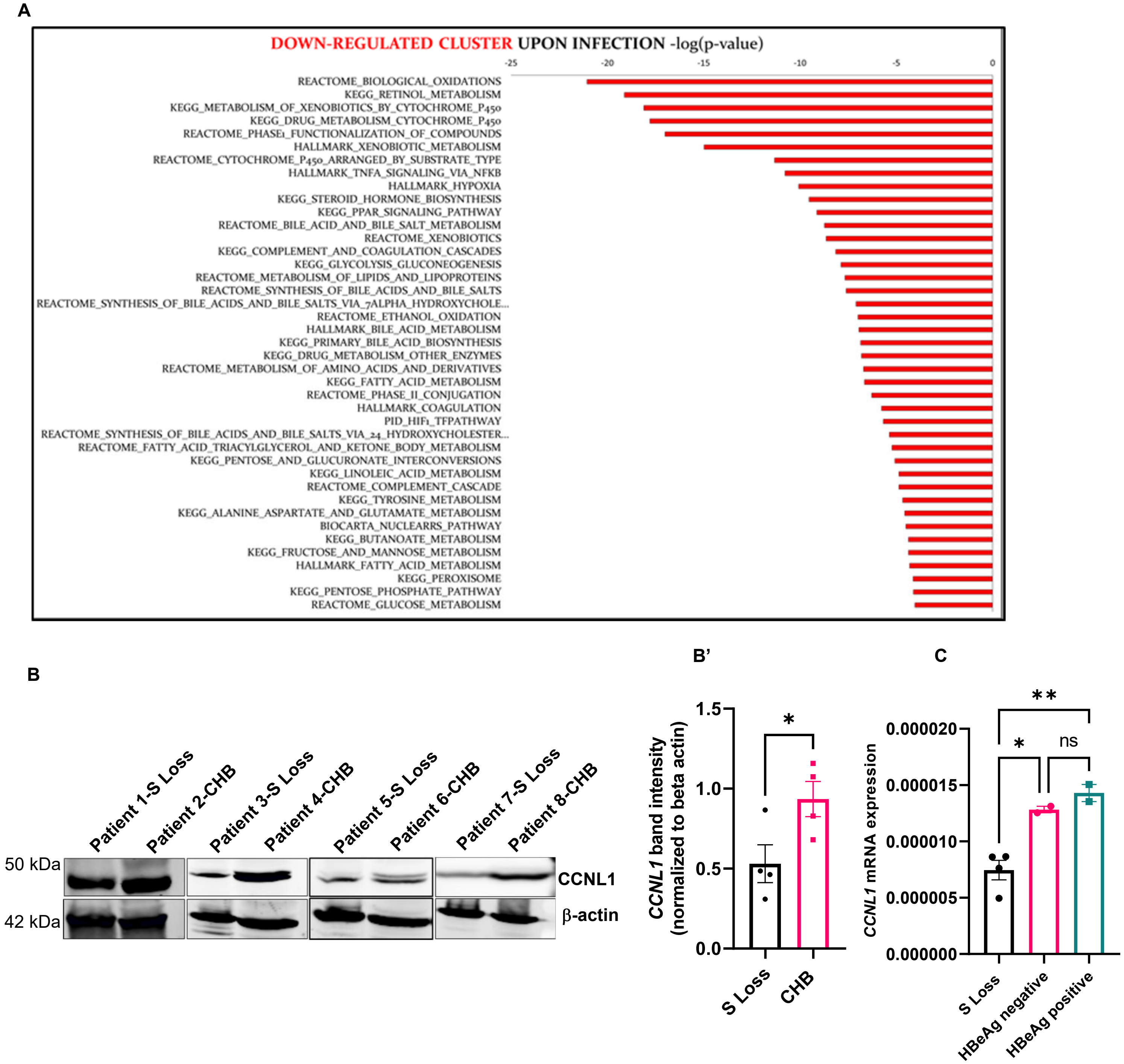

**Fig. S2 Evaluating the role of *CCNL1* during HBV infection**

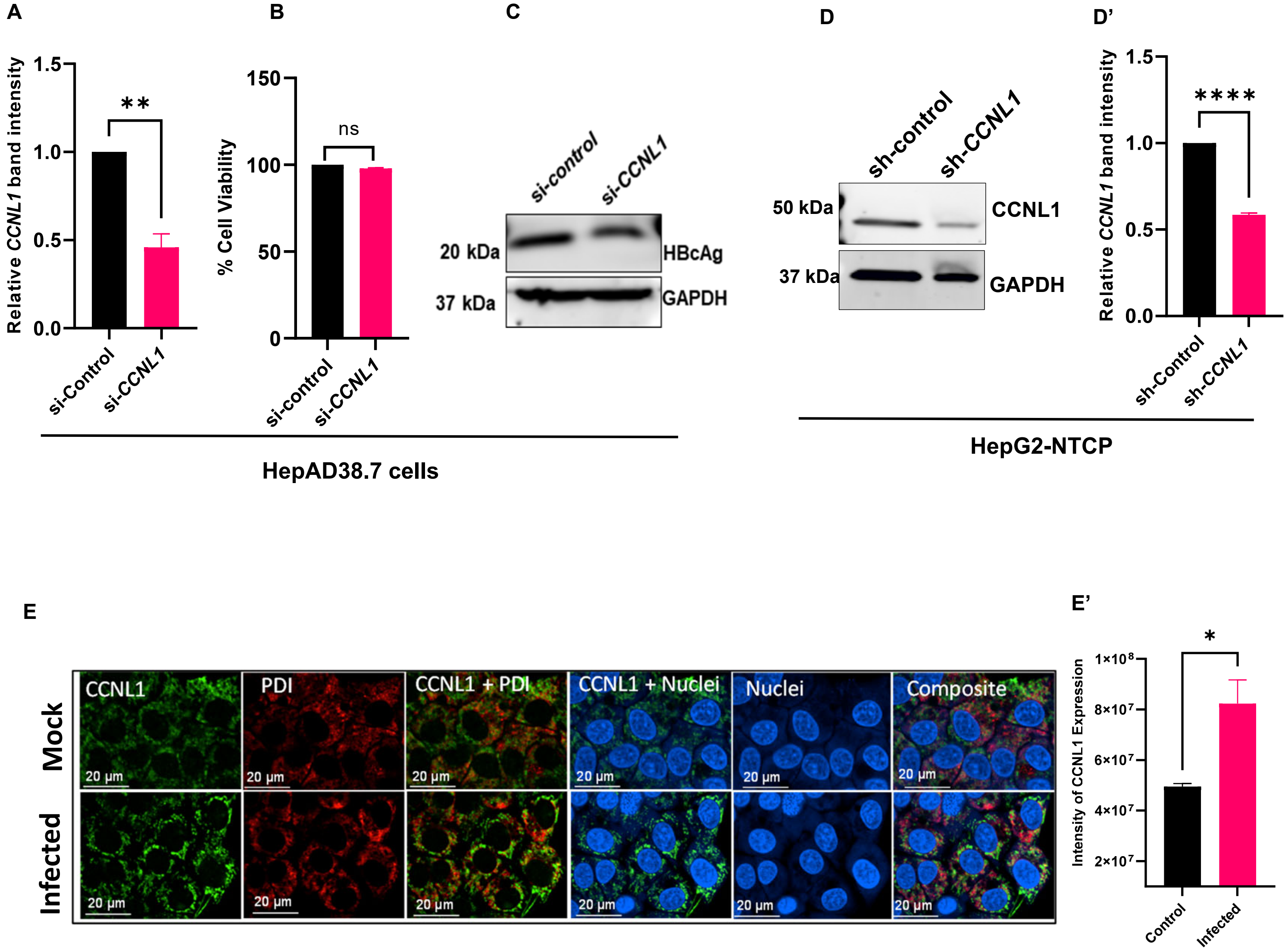

**Fig. S3 Functional validation of Cyclin L1 in PHH**

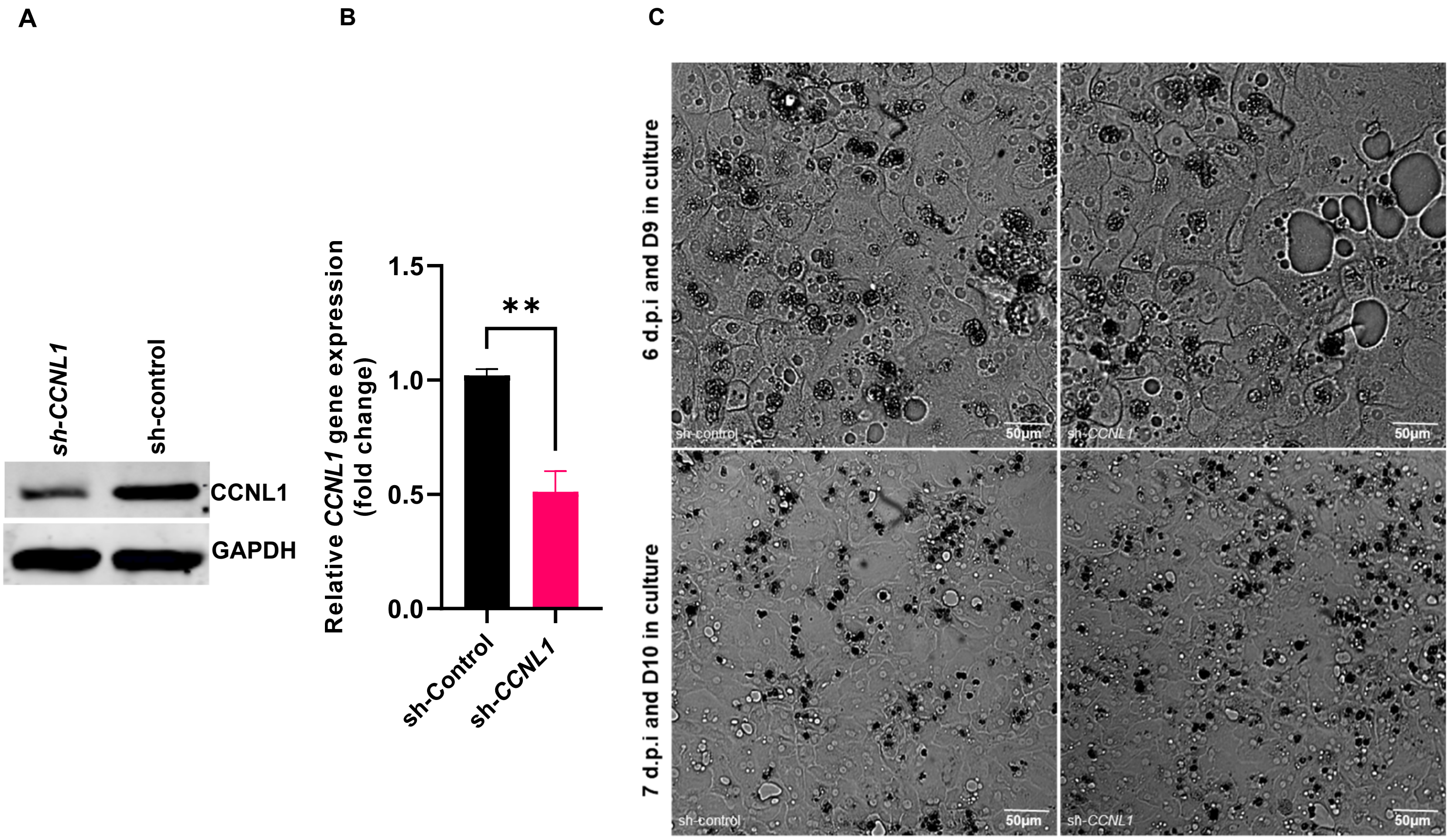

**Fig. S4 Ectopic expression of *CCNL1* in HepG2-NTCP cells enhances HBV gene expression**

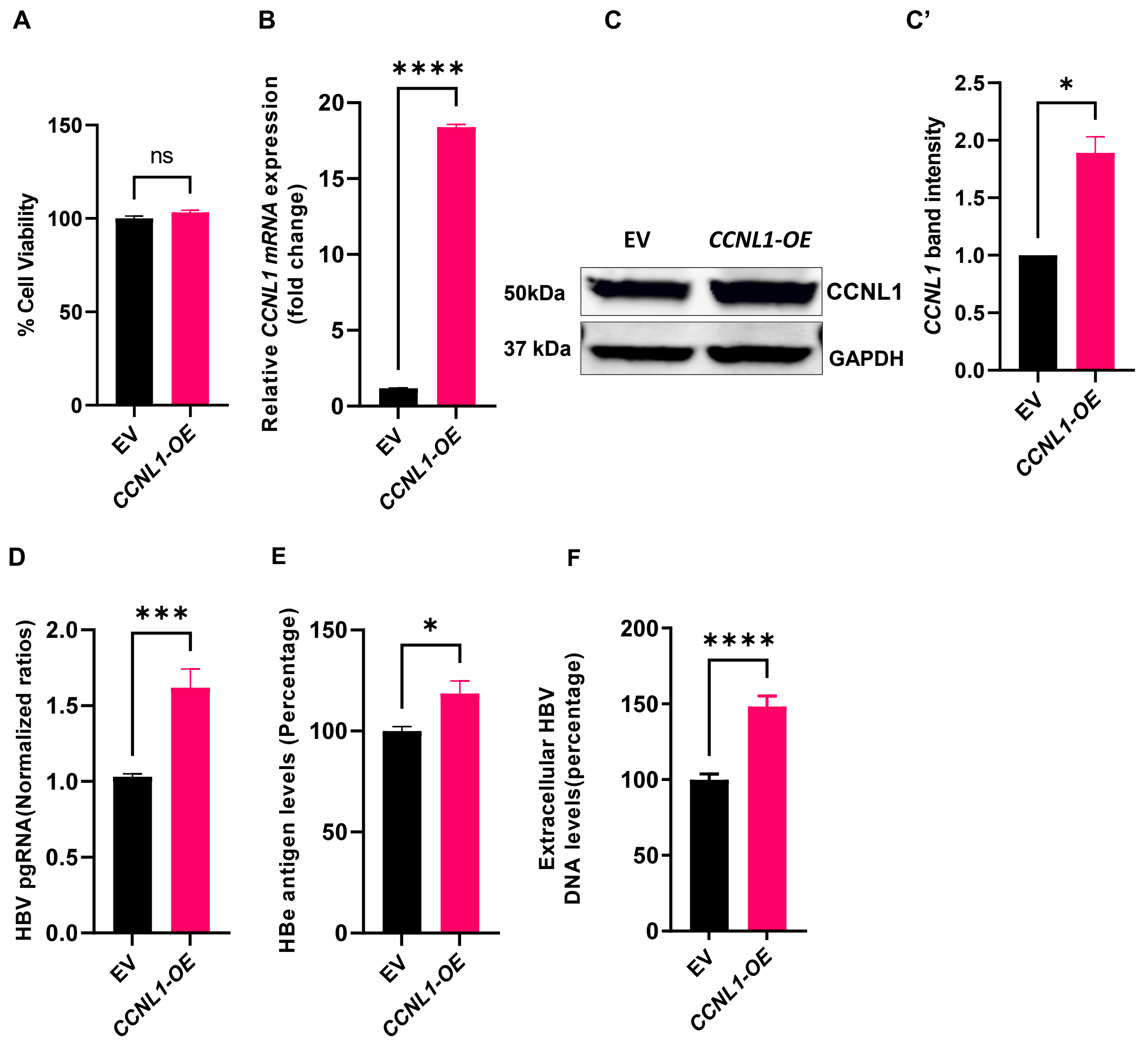

**Fig. S5: Evaluating the role of Cyclin L1 on HBV RNA splicing**

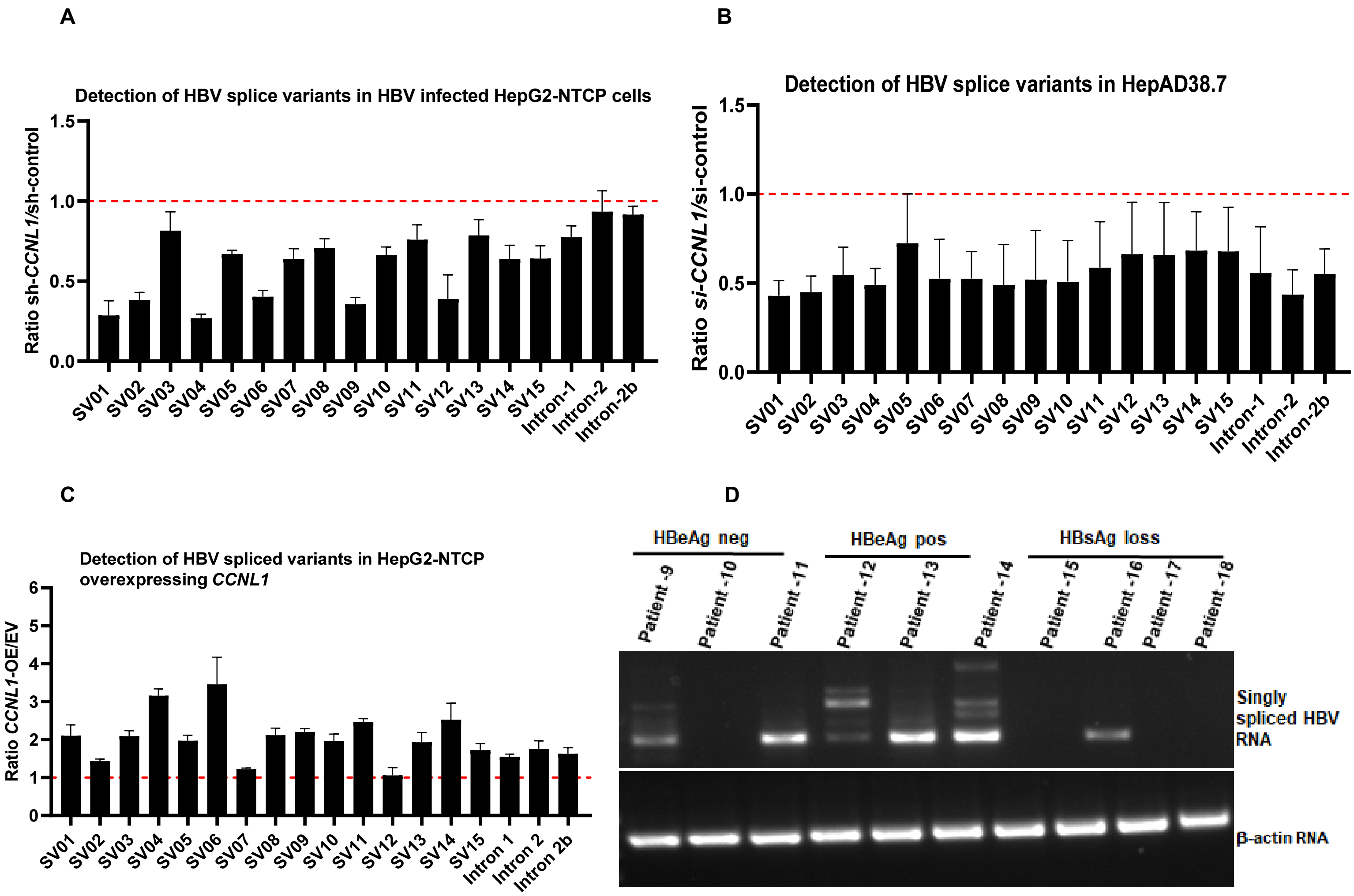

**Fig. S6 Cyclin L1 regulates HBV transcription**

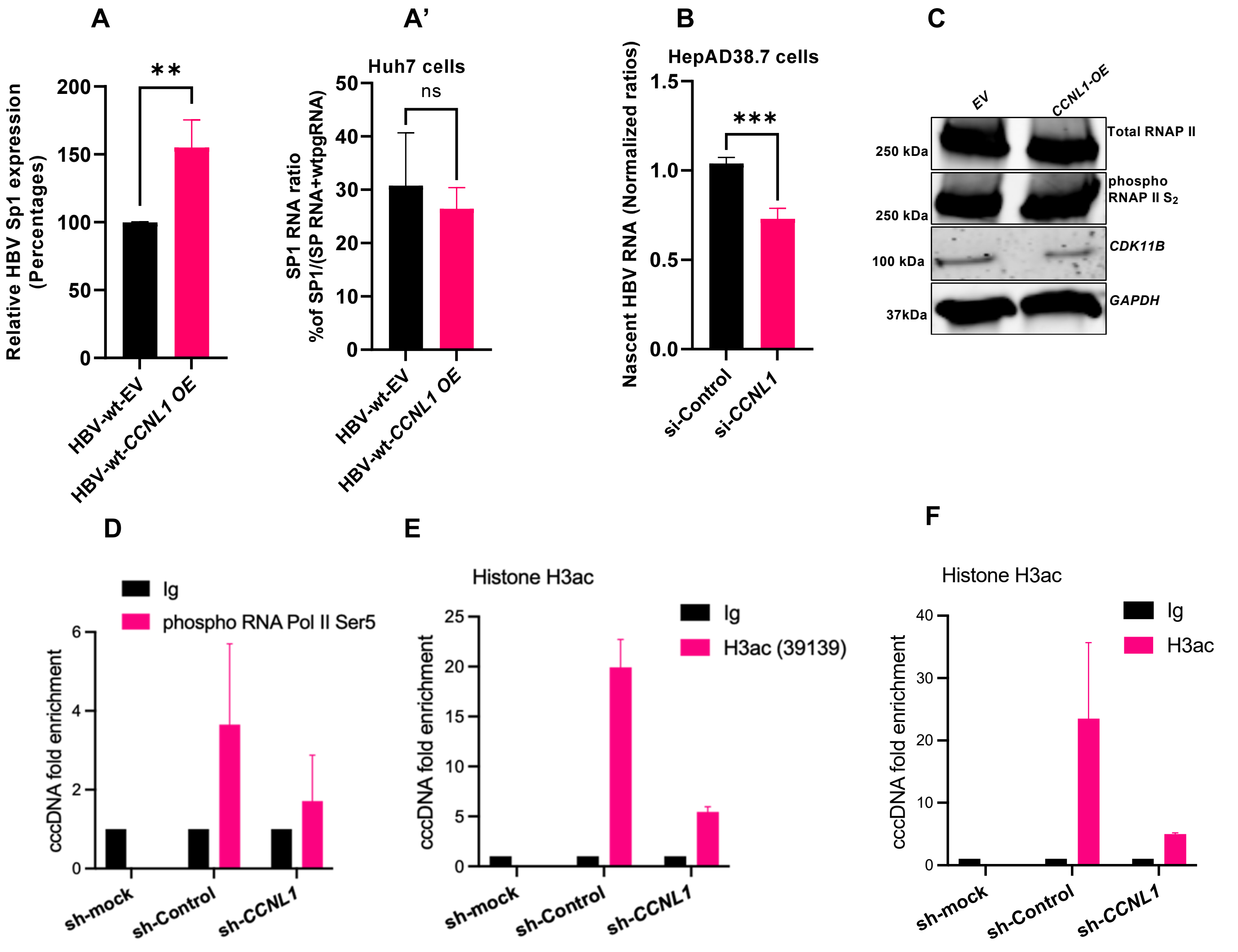

**Fig. S7 Morphometric analysis of HepG2-NTCP cells after Cell Painting and hierarchical clustering display two different clusters of sh-*CCNL1* and sh-control cells**

**A**

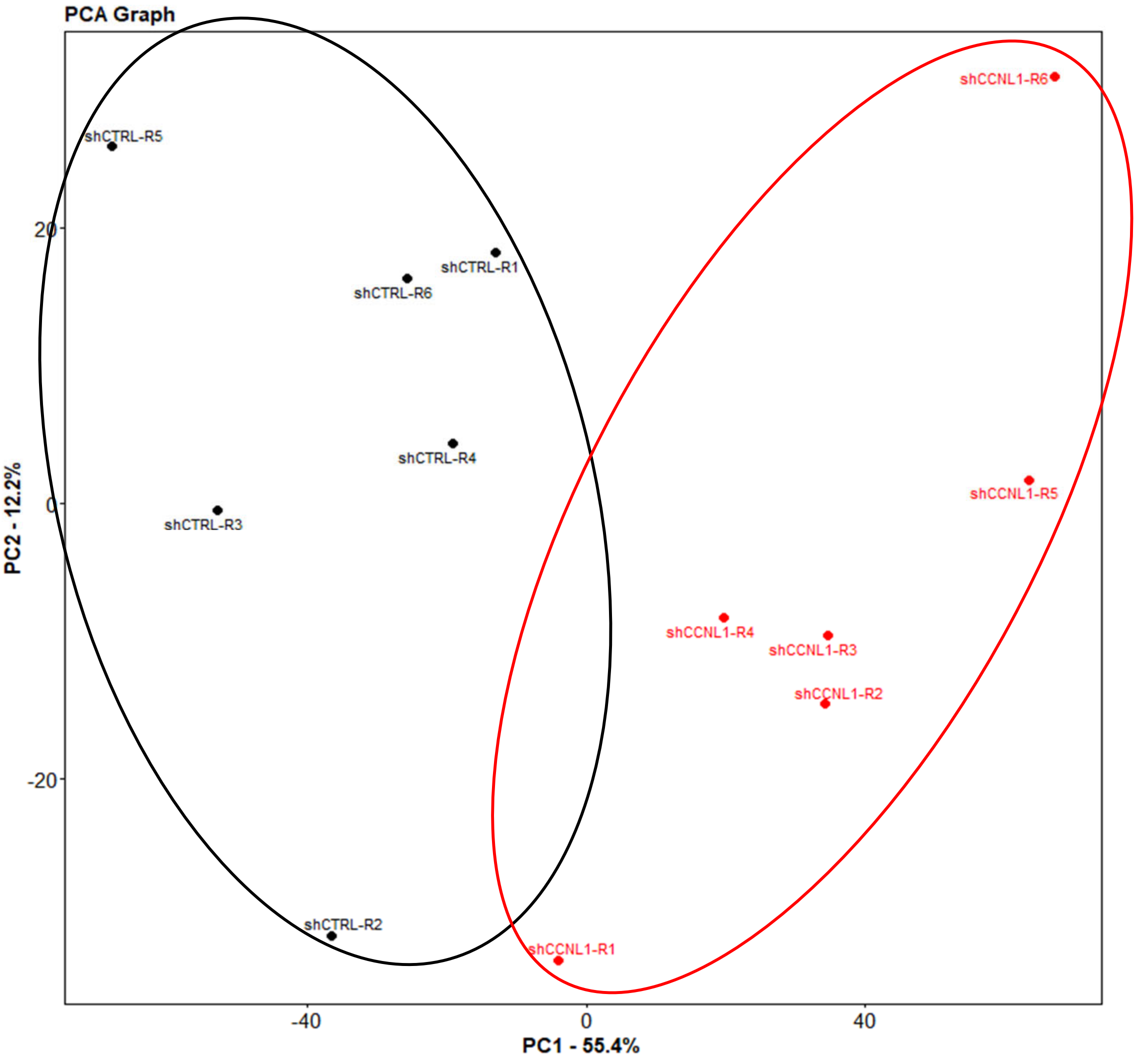
